# Dissecting splicing decisions and cell-to-cell variability with designed sequence libraries

**DOI:** 10.1101/392605

**Authors:** Martin Mikl, Amit Hamburg, Yitzhak Pilpel, Eran Segal

**Affiliations:** Department of Computer Science and Applied Mathematics; Department of Molecular Cell Biology and; Department of Molecular Genetics, Weizmann Institute of Science, Rehovot 7610001, Israel

## Abstract

Most human genes are alternatively spliced, allowing for a large expansion of the proteome.The multitude of regulatory inputs to splicing limits the potential to infer general principles from investigating native sequences. Here, we created a rationally designed library of >32,000 splicing events to dissect the complexity of splicing regulation through systematic sequence alterations. Measuring RNA and protein splice isoforms allowed us to investigate both cause and effect of splicing decisions, quantify diverse regulatory inputs and accurately predict (R2=0.75–0.85) isoform ratios from sequence and secondary structure. By profiling individual cells, we measure the cell-to-cell variability of splicing decisions and show that it can be encoded in the DNA and influenced by regulatory inputs, opening the door for a novel,single-cell perspective on splicing regulation.

## Introduction

An alternative splicing event can be the decision whether an exon is included in the mRNA (“cassette exons”), whether an intron is retained (“retained ¡ntrons”) or which of two alternative donor or acceptor sites is being used (“tandem splice sites”). The fundamental differences between these types entail peculiarities in the mode of regulation. Splicing has been shown to be influenced by a multitude of factors, ranging from local sequence motifs like poly-pyrimidine tract and branch point to epigenetic modifications. RNA binding protein (RBP) binding sites (Jangi and Sharp, 2014), DNA methylation (Lev Maor et al., 2015), secondary RNA structure (McManus and Graveley, 2011), among others, have been implicated in affecting the splicing decision, which makes disentangling the individual contributions and attributing specific functions to these regulatory mechanisms an extremely complex task when investigating native sequence contexts.

Despite extensive research, our understanding of the rules by which sequence determines splicing decisions is limited and to a large extent qualitative in nature. Approaches up to now have mainly relied on RNA sequencing data to build a computational model for alternative splicing (Barash et al., 2010; Xiong et al., 2015) or tested the effect of short randomized sequences on a nearby constant splicing event (Ke et al., 2011; Rosenberg et al., 2015). This has led to a model predicting the direction of change in alternative splicing between different tissues (Barash et al., 2010; Xiong et al., 2015) or the effect of specific point mutations on splicing (Ke et al., 2018; Rosenberg et al., 2015) or identified k-mers influencing selection of specific splice sites (Ke et al., 2011; Rosenberg et al., 2015). However, a comprehensive approach elucidating design principles of different types of alternative splicing in a context-independent manner, utilizing the power of a massively parallel reporter assay, while maintaining expression from a native locus, is still missing. Moreover, investigating splicing regulation is typically based on analyzing relative RNA isoform abundances, disregarding the downstream consequences splicing decisions can have on expression of the corresponding protein isoforms as well as their cell-to-cell variability.

Here we used a combined experimental and computational approach to unravel principles of alternative splicing in a comprehensive and quantitative way. Our approach tests rationally designed sequences and controls the genomic environment by site-specific integration, thereby reducing the regulatory complexity and enabling us to pinpoint causative sequence changes. We constructed libraries of altogether 32,789 splice site sequences and measured the effect of targeted sequence manipulations on the ratio between splice isoforms, enabling us to address many of the gaps that exist in our understanding of splicing regulation and elucidate regulatory design principles of splice sites. We follow splicing decisions in individual cells until the final gene product, allowing for a comprehensive view of splicing regulation in light of its downstream consequences and a systematic investigation of cell-to-cell variability in alternative splicing that will help to decipher the rules shaping noise in splicing decision and its functional implications.

## Results

### High-throughput testing of rationally designed splice site variants

We designed four synthetic libraries of 8551, 9608, 7473 and 7157 oligonucleotides, comprising library-specific common primers, a unique barcode and a 147–162 nt long variable region (Fig S1A). The variable region either contained (a) a retained intron flanked by exonic sequences, (b) a cassette exon flanked by intronic sequences, (c) two alternative tandem 5’ splice sites or (d) two alternative tandem 3’ splice sites. Sets of 38, 134, 81 and 96 native splice site contexts were used as basis for systematic sequence manipulations. We cloned the synthesized libraries (Agilent) between mCherry and GFP - with or without additional constant regions, depending on the splicing type - and introduced this construct in the AAVS1 locus in the human K562 cell line using zinc finger nucleases, such that every cell has one splicing reporter construct from the library and all the variants have the same genomic environment (Methods, Fig 1AB, Fig S1A). We sorted the mCherry-positive population corresponding to a single integration of the reporter transgene using flow cytometry and collected cells for RNA isolation followed by targeted RNA sequencing to quantify the abundance of different splice isoforms. We report the splicing outcome as the log-ratio between the two isoforms (Fig. 1B), i.e. spliced vs. unspliced for retained introns, included vs. excluded for cassette exons, second vs. first splice site for tandem 5’ and 3’ splice sites, as this provides a meaningful measure across all splicing types assayed here and results in a large dynamic range for quantifying effects on isoform ratio.

**Figure 1.**
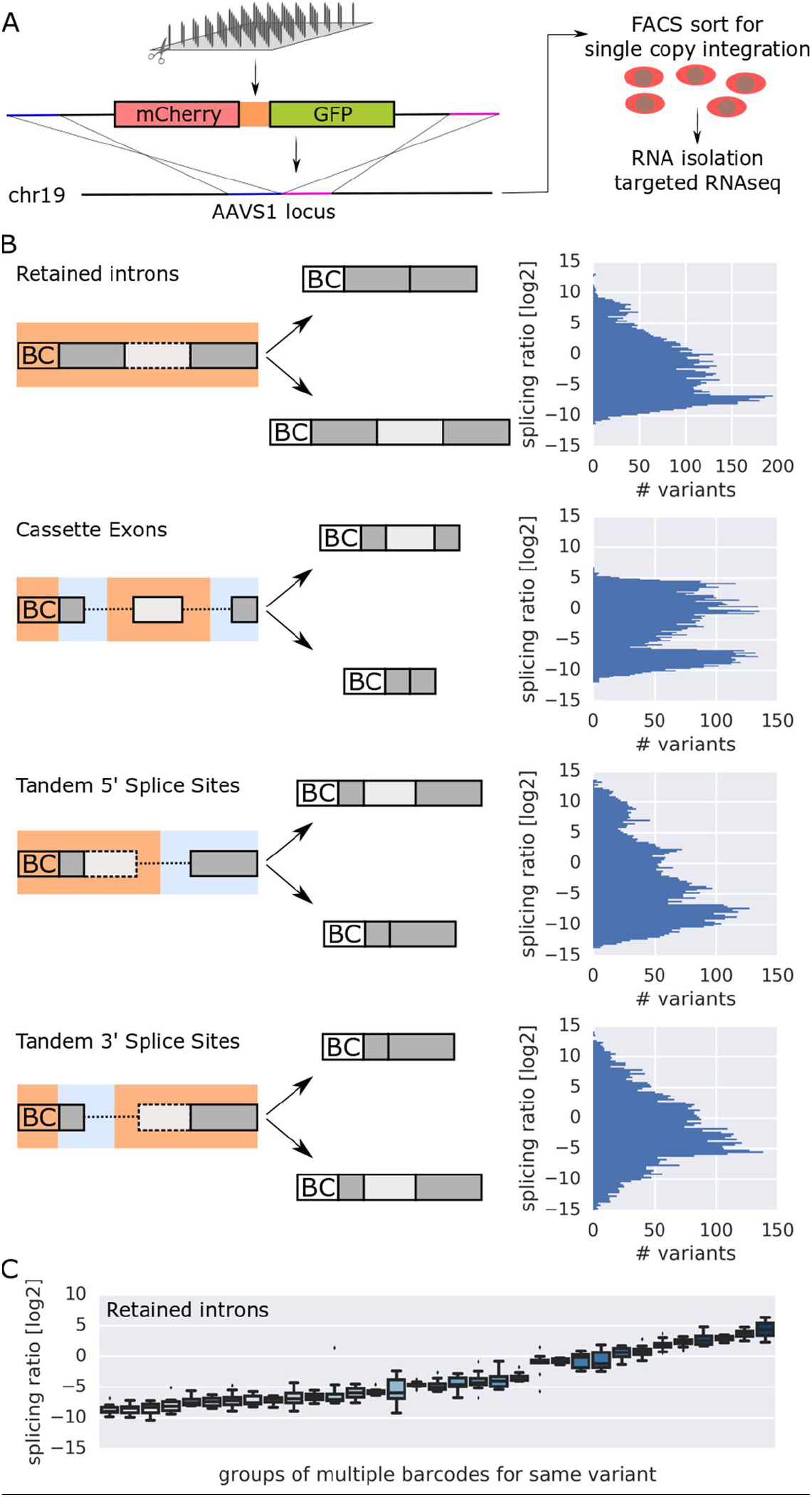
High-throughput testing of rationally designed splice site variants. A. Outline of the experimental pipeline. B. Structure of the variable (orange) and constant (light blue) regions in the four splicing type libraries, with the two splicing outcomes and the corresponding histograms. Splicing ratios close to the upper or lower limit represent predominance of the upper or lower isoform in the schematic, respectively (BC: barcode). C. Barcode controls for retained intron splicing ratios, boxplots for groups of multiple barcodes for the same sequence variant (n>7 for all groups), plotted according to their mean splicing ratio.

We confirmed the low technical noise of our system by comparing replicates (Fig S1A) and by examining groups of at least 8 independent variants with identical sequences except for the DNA barcode (Fig 1C, Fig S1B), showing that we can quantify the effect of sequence variations on splicing over a wide dynamic range.

### Splice site choice can be efficiently biased by minimal sequence changes, with tandem 5’ splice sites following a ‘first come-first served’ principle

To determine how efficiently we can bias splicing across diverse sequence contexts by manipulating only the immediate splice site sequences, we replaced the region around the endogenous splice junctions (−3 to +6 nucleotides for donor and –15 to +3 nucleotides for acceptor sites) in dozens of native sequence contexts, respectively, with consensus splice sites or non-spliceable sequences. Consensus splice sites led to efficient exon inclusion, irrespective of the native sequence context (Fig 2A), with intron-initial GT being more effective than GC (p= 1.5×10^−4^, Wilcoxon signed-rank test). Introducing an optimal branch point sequence led to a moderate increase in splice site usage in retained introns and cassette exons (Fig S2AB), but did not generally affect the choice between tandem acceptor sites (Fig S2C).

**Figure 2.**
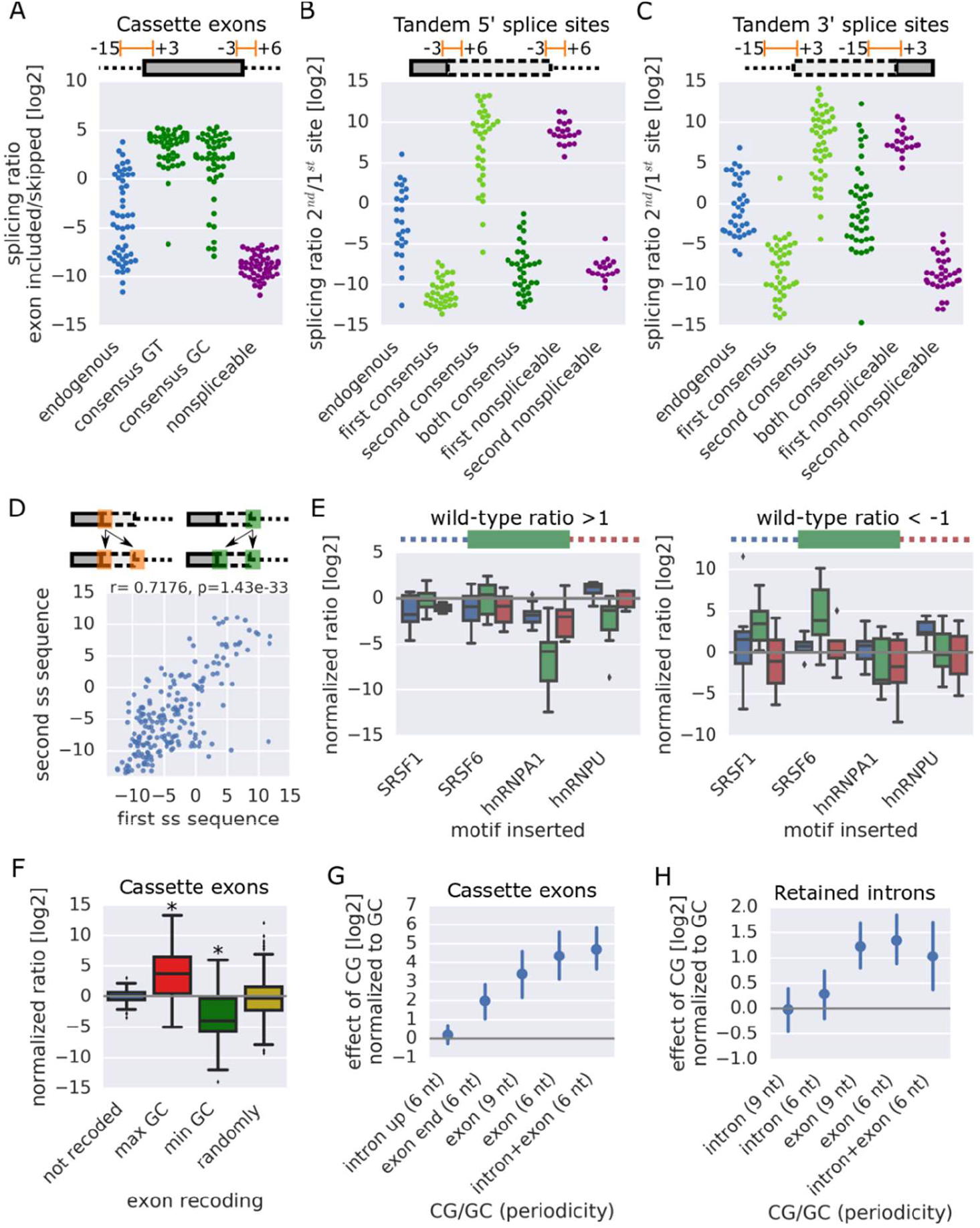
Splice site choice can be efficiently biased by minimal sequence changes (legend on next page). A-C. The indicated splice site mutations were introduced at donor (−3 to +6) and acceptor (−15 to +3) splice sites of cassette exons (A) or, in a combinatorial fashion, at one or both of tandem 5’ (B) or 3’ (C) splice sites (Methods). Data points denote splicing ratios of individual variants within the indicated group (n=17–107 for individual groups). D. Splicing ratio (2^nd^/1^st^) of tandem 5’ splice site variants in which the first splice site sequence was copied to both, plotted against the splicing ratio of variants from the same context with the second splice site sequence copied to both; each data point constitutes one endogenous context with duplicated splice site sequences of varying length (n=87). E. Distributions of the effect on the endogenous splicing ratio (=normalized ratio) of introducing a motif for the indicated splicing factor within a given region (blue: upstream intron, green: cassette exon, red: downstream intron) for contexts with a tendency for exon inclusion (left, wild-type splicing ratio >1) or skipping (right, wild-type splicing ratio <−1). Several points of insertion within this region are treated as one set (n=9–20 for each set). F. Distribution of normalized ratios (to the respective wild-type control) for recoding of cassette exons (n=99, 83, 89 and 292 variants). G-H. Mean and 95% CI of normalized ratios (to the respective wild-type control) for cassette exons (G) and retained introns (H) in which CG or GC was introduced at the indicated frequency either in the exonic or intronic regions or both (n=17–24 and n=40–58 variants in each group).

In contexts containing tandem 5’ or 3’ splice sites a consensus sequence at the first or second site led to the expected decrease and increase in splicing ratios (2^nd^/1^st^ site), respectively (Fig 2BC). When both 5’ splice sites were replaced with the consensus sequence, the first one was predominantly used across contexts (Fig 2B; p=3.3×10^−5^, Wilcoxon signed-rank test), resolving earlier conflicting evidence (Hicks et al., 2010, Rosenberg et al., 2015). For tandem 3’ splice sites no such preference could be observed (Fig 2C), indicating that the order on the transcript is a decisive factor in the choice between two donors, but not between two acceptor splice sites. This pattern held true across all potential splice site sequences (Fig S2D). Duplicating the first or the second splice site sequence showed high correlation (Fig 2D), suggesting that the context-specific contribution to splice site choice is set by sequence properties away from the immediate splice site.

To quantify the effect of additional regulatory elements, we introduced binding sites for common splicing factors and splicing-regulatory sequence motifs from previous studies (Ke et al., 2011, Rosenberg et al., 2015) at different positions in dozens of native contexts and tested for their influence on donor and acceptor sites. Motifs often showed location- and splicing type-dependent activity (Fig S2E). Single binding sites could have dramatic effects on exon inclusion levels (Fig 2E). Their activity often depended on the “quality” of the splice site (e.g. hnRNPA1, SRSF6), but less on the precise binding site location (Fig S2F), and was affected by coinsertion of other binding sites (e.g. hnRNPA1 +SRSF1; Fig S2G).

### GC content in general and CG dinucleotides in particular strongly affect splicing decisions

Introns and exons differ in their GC content (Amit et al., 2012). To assess the potential for regulation based on GC content alone, given a desired protein outcome, we recoded native splice site regions and measured the effect on splicing. Recoding of a cassette exon for highest or lowest possible GC content had strong opposite effects of similar magnitude (mean fold change ~2^4^, Fig 2F), indicating that endogenous cassette exons tend to be not committed to either high or low splicing efficiency based on their GC content alone, leaving a lot of regulatory potential to influence splicing in either direction.

DNA methylation at the cysteine residue in CG dinucleotides has been proposed as another means for regulating splicing (Lev Maor et al., 2015). We introduced CG or GC at different frequencies in 30 cassette exons and quantified the effect conferred by introduction of CG, which is potentially methylated, as opposed to GC (Fig 2G). When introduced in the exon, CG biased splicing towards inclusion of the exon (p=1.0×10^−22^, Wilcoxon signed-rank test) in a dose-dependent manner. Having additional CGs in the intron did not interfere with this positive effect (p=0.22), suggesting that the presence of CGs increases usage of an already defined cassette exon, as opposed to making it distinguishable from the surrounding intronic region.

Changing the GC content in and around retained introns had only the potential to repress, but not to enhance splicing (Fig S2H). A differential effect of CG vs GC dinucleotides in the exon could be observed also here (Fig 2H; p=9.0×10^−11^, Wilcoxon signed-rank test), supporting recent observations showing a relationship between loss of DNA methylation and increased intron retention (Kim et al., 2018; Wong et al., 2017).

### Coordinated and antagonistic effects of exons and introns shape the overall splicing outcome

To assess the potential of individual building blocks to confer regulatory properties from one context to another, we substituted exonic and intronic components of alternative splice sites with sequences from native splice sites without evidence for alternative splicing (Fig 3A). While the constitutive effect of 3’ ends of introns can generally be conferred to other contexts (Fig S3A), this is not the case for sequences surrounding the donor splice site (Fig 3B). Effects for pairs of exon and intron sequences taken from the same native context showed negative correlation (Fig 3C), hinting at antagonism between exonic and intronic components.

**Figure 3.**
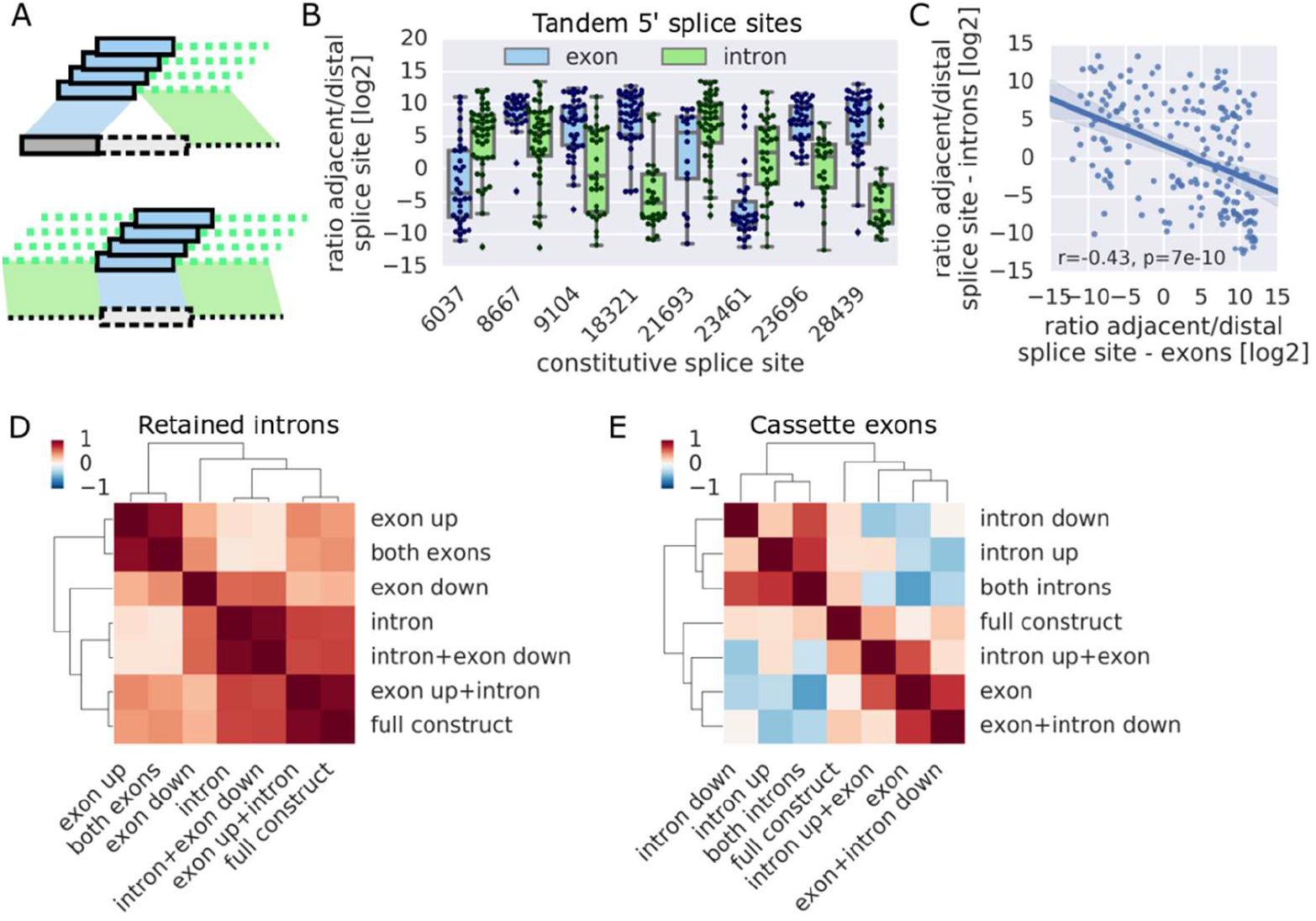
Antagonistic interactions between building blocks shape the overall splicing outcome. A. Schematic for the replacement of components of the alternative splice sites tested with elements from a set of “constitutive” splice site sequences, shown for tandem 5’ splice sites and cassette exons. B. Distribution of log-ratios of usage of the adjacent in tandem 5’ splice sites in variants with the upstream exon (blue) or the downstream intron (green) replaced with the corresponding exonic or intronic part from a set of eight constitutive splice sites (n=17–47 in each group). C. Each datapoint represents the log-ratio of usage of the proximal vs. distal splice site relative to the replaced component (exon or intron from the same native context, n=279). D-E. Clustered heatmaps showing Pearson correlation coefficients for all pairwise combinations of effects on splicing ratio in (D) retained introns or (E) cassette exons conveyed by the indicated elements across contexts (n=96–215 and 48–143 for each combination).

To test for coordinated effects between building blocks of native splice sites, we replaced components of 38 retained introns and 30 cassette exons with sequences from 5–6 length-matched unrelated introns and exons (Fig 3DE). High correlation in associated splicing ratios could be observed between introns and their corresponding downstream exon across library contexts (Pearson r=0.57, p=2×10^−13^, Fig 3D, Fig S3B), suggesting coordinated effects in the native context. In the case of cassette exons, intronic and exonic elements showed strong negative correlation (r=−0.6, p=4×10^−7^, Fig 3E, Fig S3C), arguing for antagonistic effects creating a balance between components favoring or disfavoring splicing that gives rise to the endogenous splicing decision.

### Accurate prediction of splicing ratios based on sequence and RNA structure

Having measurements for large collections of splice site variants from a constant genomic environment, we wanted to undertake a task that has proven challenging in the field of splicing: To quantitatively predict splicing based on sequence. We used splice site strength (as determined using MaxEntScan (Yeo and Burge, 2004)), hexamer counts, cumulative binding scores for 160 RNA binding proteins (ATtRACT, Giudice et al., 2016) and RNA secondary structure, alone and in combination, as features and trained a Gradient Boosting Regressor (Methods; Fig 4, Fig S4A), allowing us to quantitatively predict splicing ratios of unseen variants for all splicing types with high accuracy (R^2^ between 0.774 and 0.825) and recovering known splice site components in an unbiased way (Fig S4B).

**Figure 4.**
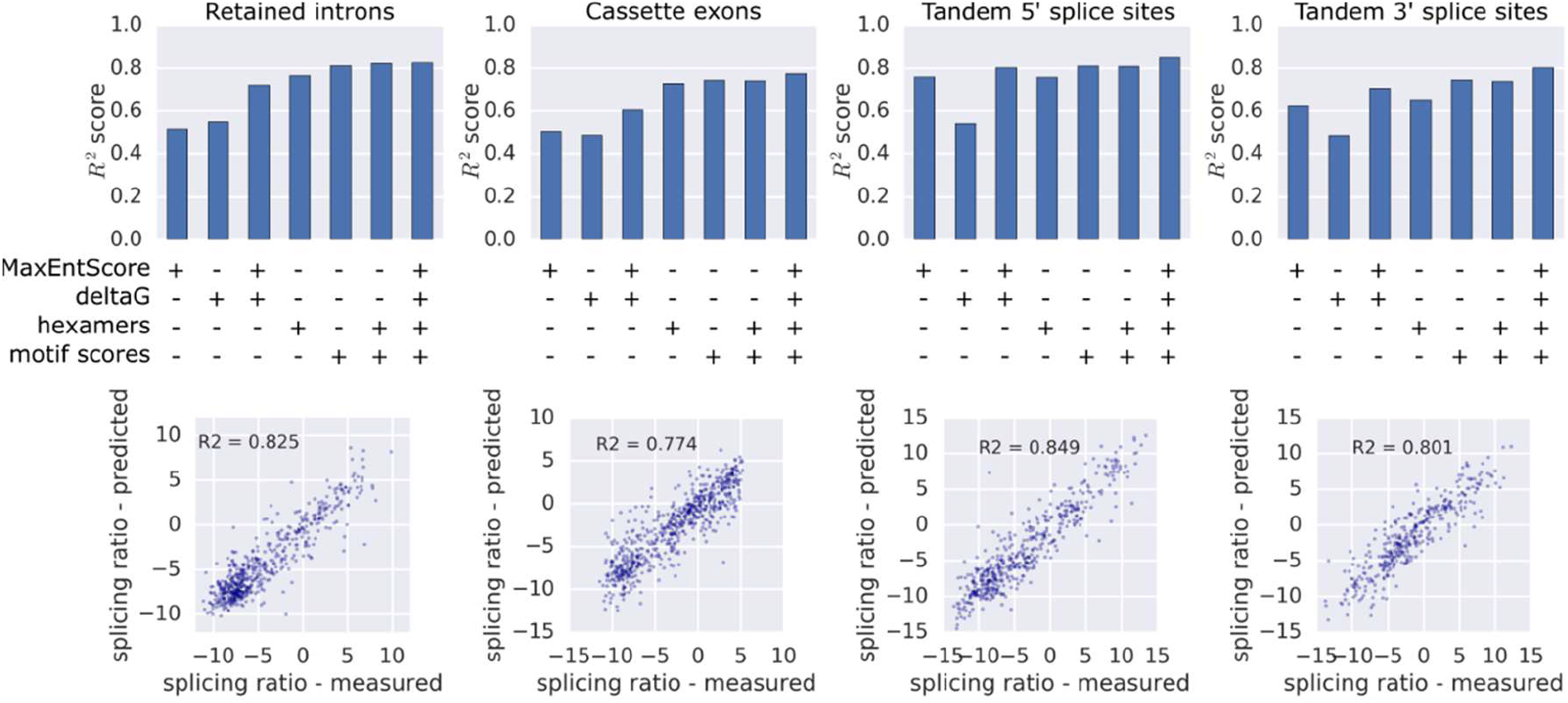
Accurate prediction of splicing ratios. R^2^ scores for predicting splicing ratios of a set of variants not used at any point to build the model (10% of library variants) by Gradient Boosting Regression with the indicated sets of features (top) and measured splicing ratios vs. predictions based on a combination of hexamer counts, RBP motif scores, MaxEntScan scores and secondary structure features (bottom) for the indicated splicing types.

Using only the predicted minimum free energy (ViennaRNA) of regions around splice sites, we achieved R^2^ scores between 0.48 and 0.55 (Fig 4), indicating that secondary structure alone, without any sequence information, can be predictive of splicing outcome. This is in line with the strong effect introducing a hairpin around or downstream of splice sites has on splicing (Fig S4C) and the pronounced preferences for secondary structure (Fig S4D), e.g. for the pyrimidine tract to be paired in retained introns, but unpaired in alternative 3’ splice sites, reflecting fundamental differences in processing of different splicing types.

### Splicing readout at the protein level reveals differential posttranscriptional fates

To be able to follow splicing decisions in individual cells to the final gene product, we constructed our library in a way that allows us to quantify splicing with a bifluorescent reporter (Gurskaya et al., 2012) in large scale (Fig 5A). Only if the intron is removed is the downstream *gfp* in frame and are both mCherry and GFP made into protein. The ratio of GFP vs. mCherry fluorescence is a sensitive measure of protein isoform ratios in individual cells.

We sorted the pool of cells, each carrying one variant, into 16 bins according to their GFP/mCherry ratio and sequenced genomic DNA from all the bins to unravel the distribution of each variant (Fig 5A), which provides a measure of both the population average as well as the variability of splicing decisions at the single-cell level.

We previously demonstrated that similar approaches are highly accurate and reproducible (Weingarten-Gabbay et al., 2017, Vainberg Slutskin et al., 2018). Results for groups of identical barcodes (Fig S5A) and the associated bin profiles (Fig S5B) corroborate the low technical noise we are able to achieve.

RNA- and protein-based readouts are well correlated (Pearson r=0.75; Fig 5BC), but show particular differences. RNA expression levels increase with efficiency of intron removal (Fig 5CD), but protein expression levels are equally high at low and high splicing values (Fig 5F), suggesting that – on average – transcript variants lacking clear splicing signals can yield similar translational outputs as efficiently spliced variants. No such discrepancy could be observed for alternative 5’ splice sites (Fig 5EG), where the decision is not whether to splice or not, but which splice site to use. Variants with intermediate splicing levels are more likely to be degraded due to failed processing, resulting in both lower RNA and protein levels.

Relative intronic GC content is negatively correlated with the RNA, but not the protein splicing ratio (Fig 5C), indicating that influences of GC content on splicing efficiency are buffered at the protein level. To identify specific sequence features mediating the discrepancy between RNA and protein we computed the difference in Pearson correlation coefficients between all intronic hexamer counts and either RNA- or protein-based splicing readouts (Fig S5C). Several properties of optimal introns (low GC content (Fig S5D; see also above), the consensus 5’ splice site GTAAGT and pyrimidine-rich features (Fig S5C)) were among the hexamers with the largest negative impact on protein-based splicing values.

**Figure 5.**
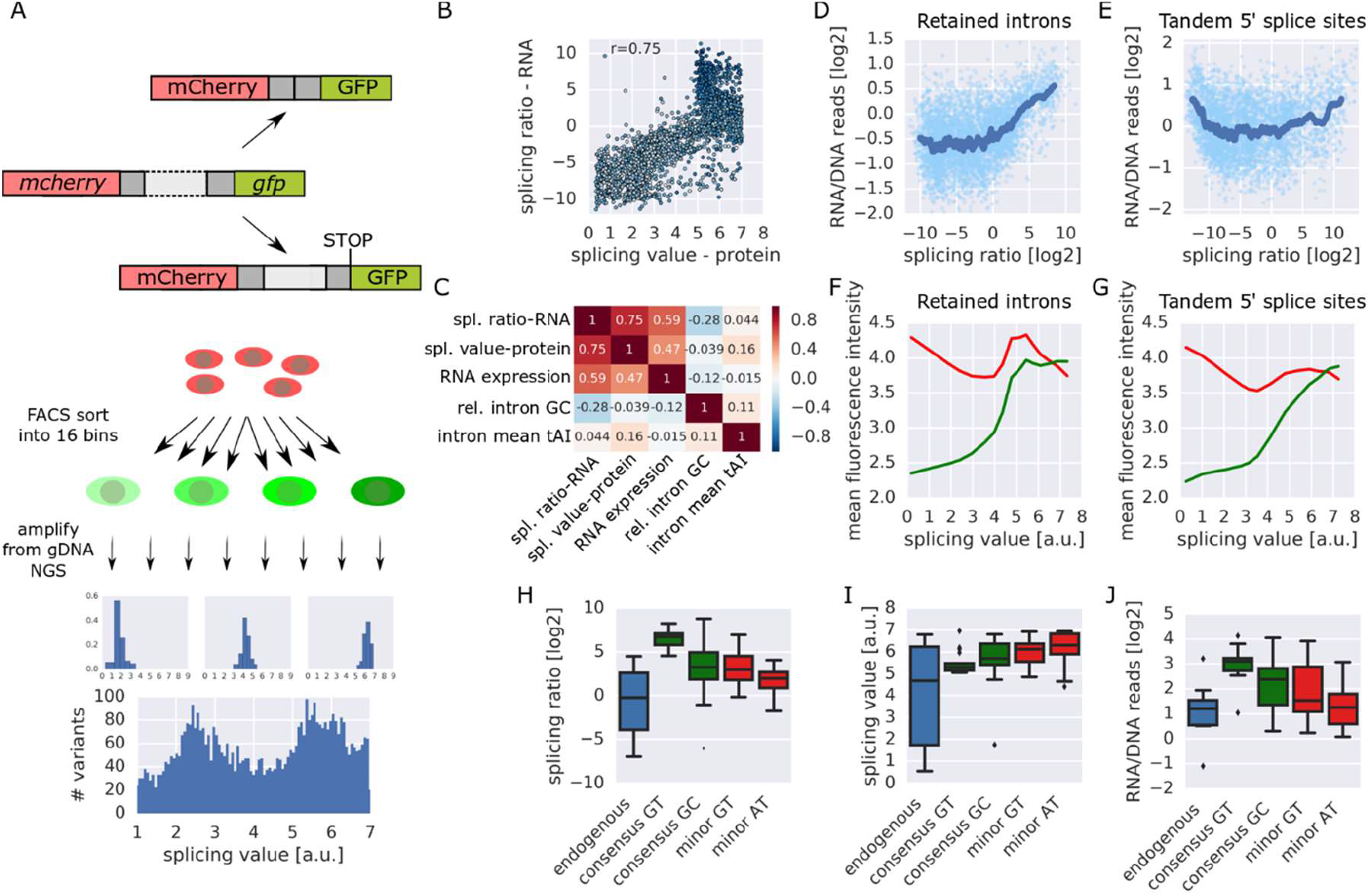
Quantifying protein isoform ratios reveals differential posttranscriptional fates. A. Outline of the experimental pipeline for obtaining protein-based splicing measurements for retained introns. B. RNA-based splicing ratios plotted against protein-based splicing values for the retained intron library. C. Pearson correlation coefficients between RNA-based splicing ratios, protein-based splicing values, RNA expression levels (log ratio of RNA/DNA reads), intronic GC content relative to the surrounding exons and mean tAI of retained introns. D-E. Log ratios of RNA/DNA reads (=RNA expression levels) plotted against splicing ratios for the retained intron (D) and tandem 5’ splice sites (E) library. F-G. Mean mCherry and GFP fluorescence intensity for cells from the retained intron (F) or tandem 5’ splice sites library (G) sorted into each of the 16 bins are plotted against the respective splicing value (i.e. the median log ratio of GFP/mCherry fluorescence intensity). H-J. Box plots show the distribution of RNA-based splicing ratios (H), protein-based splicing values (I) and log ratios of RNA/DNA reads (J) of individual variants within the indicated group (n=22–59).

We therefore compared 38 native retained introns in which the immediate splice sites were replaced with either the consensus sequences for the major spliceosome or less efficiently processed alternatives (5’-GC, minor spliceosome). While a consensus for the major spliceosome resulted in higher ratios of spliced/unspliced transcripts (Fig 5H), corresponding variants with a 5’-GC or minor spliceosome-specific sequences were associated with a higher fraction of the spliced, GFP-containing isoform on the protein level (Fig 5I).

Mean GFP intensity reaches a plateau around splicing values corresponding to processing of consensus sites by the major spliceosome (splicing value ~5; Fig 5F). Higher GFP/mCherry ratios appear to be due to lower expression levels of the mCherry-only protein product, indicating that the observed discrepancy is due to reduced translational output from the unspliced isoform. RNA expression levels are strongly influenced by the efficiency of splicing and the machinery involved. GT-consensus sequences lead to dramatically increased RNA levels (Fig 5J, p=0.0026, Wilcoxon signed-rank test), likely exceeding the capacity of downstream processing steps.

Our observations might reflect a bottleneck due to inefficient processing. Here, a larger proportion of RNA molecules than in the case of the optimal consensus would not be processed immediately after transcription. This unspliced pre-mRNA is not exported and translated, but is detectable on the RNA level. Our results therefore provide evidence that less efficient splicing can yield more clear-cut choices between protein isoforms than maximal splicing efficiency.

### Splicing noise can be encoded in the sequence and affected by regulatory inputs

Evidence from transcriptome-wide approaches (Marinov et al., 2014; Shalek et al., 2013) and studies focusing on individual genes (Gurskaya et al., 2012; Marinov et al., 2014) provide examples for cases where bulk splicing measurements do not adequately reflect splicing decisions on the single-cell level, with potentially far-reaching functional biological implications as observed in other areas of gene regulation (Eldar and Elowitz, 2010). Our approach allows us to quantify the variability in splicing between cells based on the distribution across bins (Fig S6A). The strength of splicing noise (variance/mean) is negatively correlated with splicing efficiency (Fig 6A). To account for this dependency we used the deviation of noise strength from a linear fit as a noise measure and refer to this as the noise residual.

**Figure 6.**
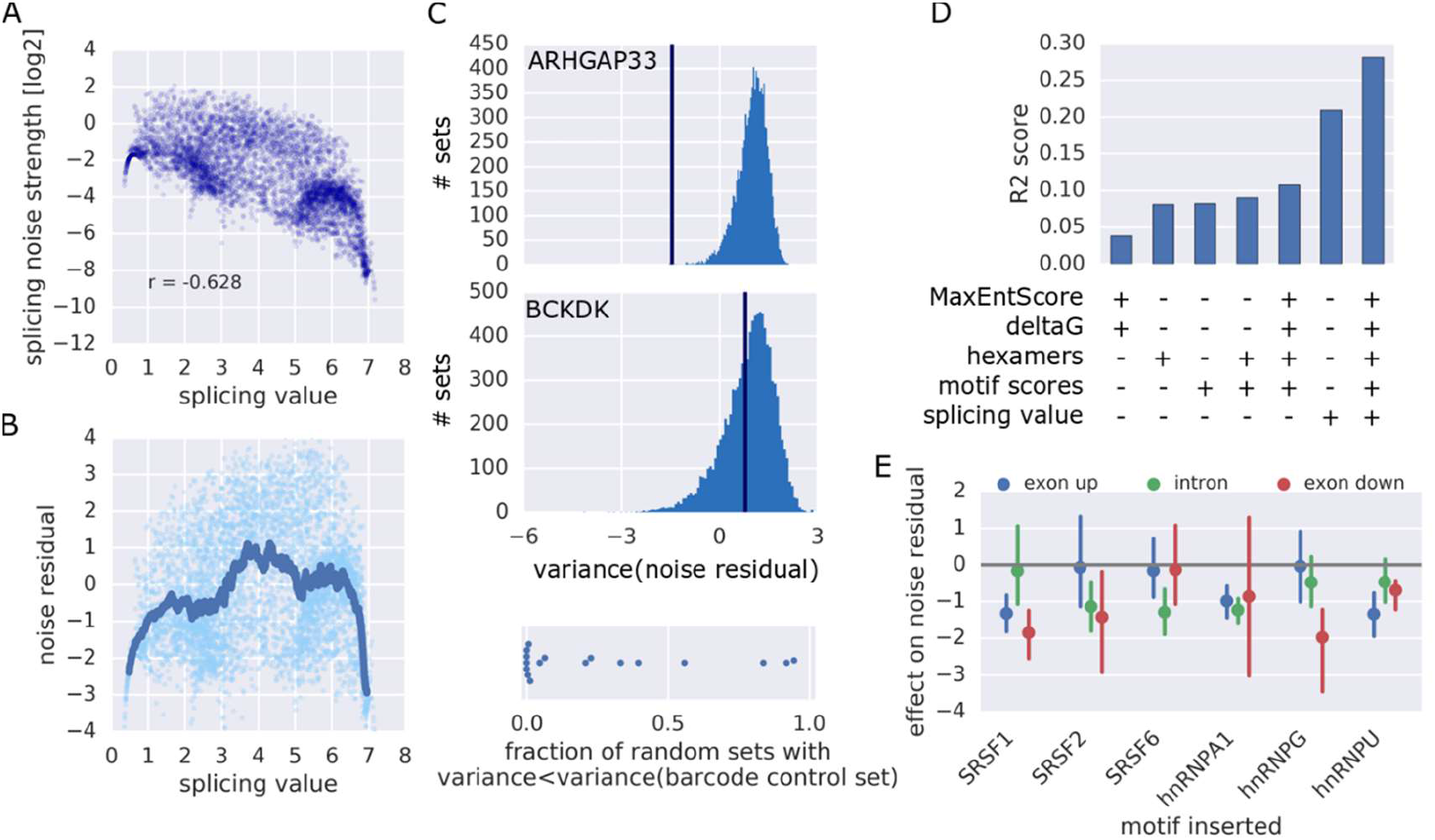
Cell-to-cell variability of alternative splicing can be encoded in the DNA sequence. A. Noise strength (variance/mean) is plotted against mean splicing values for the retained intron library. B. Scatterplot and rolling average (window size=100) showing noise residuals and splicing values. C. Representative examples for comparing the variance of noise residuals for groups of individual barcode controls to variances for 10.000 sets of equal size, randomly picked from library variants in the same range of splicing values. The vertical blue line indicates the variance of the barcode control group. The bottom panel shows the fraction of random sets with a variance smaller than the one of the corresponding barcode control set. D. R2 scores for predicting splicing noise residual using the indicated features. E. Mean and 95% CI of the effect on the noise residual of introducing a motif for the indicated splicing factor within a given region. Several points of insertion within this region are treated as one set (n=4–18 in each group).

The noise residual tends to be higher for intermediate splicing values (>3 and <5), while splicing decisions seem to be less noisy when there is one dominant isoform (<3 and >5; Fig 6B).

But can the cell-to-cell variability of splicing be determined by the DNA sequence? To test this, we used sets of at least 8 identical splice site sequences and checked if the variability within these sets is smaller than would be expected by chance. For each set, we compared the variance of noise residuals to the distribution of variances from 10.000 randomly chosen sets of splicing value-matched variants (Fig 6C). While some sequences show significantly lower variance in noise levels (especially those with intermediate splicing values; Fig. S6B), in other cases the within-group variability is as high as in randomly picked sets, indicating that noise level can be encoded in the sequence, but this is not implemented for every splicing event.

Despite this limitation we could account for over 10% of the variability with a predictive model using our set of sequence and structural features (Fig 6D, Fig S6CD). The mean splicing value remained the most informative feature and could alone explain more than 20% of the variability. Using all of the above further increased the performance of prediction to R^2^=0.281, indicating that the sequence and structural features predict noise not only through the mean splicing value, but capture independent contributions of sequence and secondary structure.

Can splicing noise be affected by regulatory inputs? Introduction of e.g. an SRSF1 binding site in the exon led to a significant reduction in noise residual (Fig. 6E, p=0.0093, Wilcoxon signed-rank test), with no significant effect on mean splicing values or expression levels (Fig S6E), showing decoupling between mean and noise. This is in line with increased variability in MKNK2 splicing observed in SRSF1 depleted cells (Waks et al., 2011). Introducing a splicing factor binding site generally led to reduced noise, which could be due to greater certainty in the splicing decision (even without affecting mean splicing values), leaving less room for cell-to-cell variability.

## Discussion

Here, we used rationally designed libraries, consisting of altogether 32,789 variants, to address fundamental questions in splicing regulation. This allowed us to dissect and compare the different regulatory inputs in a quantitative way and identify design principles of alternative splicing events, considering the process in its entirety, from the processing of the RNA to the level of the final functional gene product. Our study goes beyond previous approaches by (a) yielding readouts for RNA and protein isoform ratios and expression levels, (b) making use of a fully designed sequence library, allowing us to reduce the complexity of splicing regulation, (c) integrating each variant in the same genomic location, thereby mimicking expression from a wild-type locus, (d) surveying different splicing types in a comprehensive and comparative way, (e) testing our targeted sequence manipulations in dozens of contexts, eliminating potential biases due to specific effects of sequence changes on the one splicing event typically used in a reporter assay.

Using this approach, we can reproducibly detect even small changes in splicing ratios and quantitatively predict splicing of novel variants with high accuracy (R^2^ between 0.76 and 0.85). Our approach to elucidating the context-independent principles of splicing regulation is complementary to studies using endogenous RNA sequencing data to establish a splicing code for prediction of drastic changes in splicing behavior between cell types (Barash et al., 2010; Xiong et al., 2015).

Our results show that it is relatively straightforward to build an optimal splice site; simply using the consensus splice site sequence can efficiently trigger splicing, no matter what the surrounding sequences. Large effect sizes can be achieved with even single splicing factor binding sites, altering codon usage and introducing CG dinucleotides, demonstrating that each regulatory input by itself has the ability to significantly bias splicing in most native contexts. And yet, cells evolved to have seemingly “suboptimal” splice sites, which maximizes the potential for dynamic regulation, but can also serve to ensure optimality at the level of protein isoforms.

Splicing occurs at the RNA level, but it is typically the resulting protein products whose functional differences constitute the significance of this process. Our dual RNA- and protein-based assay revealed properties of splice sites associated with differential downstream fates of isoforms and highlighted that capturing a snapshot at the RNA level might not always reflect the consequences of an alternative splicing event at the level of the final functional gene product.

While the role of noise in other aspects of gene expression has attracted a lot of attention over the last years (Elowitz et al., 2002; Kaufmann and van Oudenaarden, 2007; Ozbudak et al., 2002), assessing the variability of splicing decisions has been lagging behind, largely due to technical limitations. Even single-cell RNAseq approaches are limited in their power to detect cell-to-cell differences between splice isoforms differing in only a couple of nucleotides. Here, we established an assay that is able to assess the cell-to-cell variability of splicing decisions in large scale by measuring the protein output of alternative isoforms. We show that the level of stochasticity can be encoded in the DNA. In general, our data present noise as a complex property of splicing events, which is in part a passive consequence of the stochastic nature of gene expression and the uncertainty associated with intermediate splicing efficiencies, but can be influenced by specific sequence elements and properties of a splice site.

## Materials and methods

### Synthetic library design

#### General design notes

Oligonucleotides were designed to maintain a constant length of 210 nt. Restriction sites used for cloning and splice site sequences apart from the assayed donor and acceptor sites were excluded from the design. All the variants were composed of an 18 nt forward primer, 12 nt barcode sequence, 162 nt variable region and 18 nt reverse primer sequences. Barcodes were designed to differ from any other barcode in the library in at least 3 nt. In the case of cassette exons and tandem 3’ splice sites, which required a subsequent cloning step and therefore additional internal restriction sites, SpeI and AatII sites were introduced after the barcode sequence with a 3 nt spacer between them, leaving 147 nt for the variable region. The unique primer sequences at the 5’ and 3’ ends were used for targeted amplification of the variants from the pool of synthesized oligonucleotides.

#### Selection of contexts

For all four libraries used here, endogenous sequence contexts (38, 134, 81 and 96 for the retained intron, cassette exon, tandem 5’ and tandem 3’ splice sites libraries, respectively) were selected based on (a) prior testing in a low throughput pilot screen in the same context or (b) by selection of suitable splice sites from publicly available RNA sequencing data for K562 cells (Encode, polyA RNA-seq of K562, Gingeras Lab, accession number ENCFF000HFA; intron length 70–118 nt, exon length 23–89 nt, distance between tandem 5’ splice sites 2–77 nt, distance between tandem 3’ splice sites 2–59 nt). Contexts were chosen such that a wide range of putative splicing ratios would be covered and the alternative splicing event would lead to isoforms with a different downstream reading frame used. In the case of retained introns, a frame shift was introduced in the intron unless intron retention already led to one or the intron contained a stop codon, in order to allow for discrimination of the isoforms on the protein level based on GFP being in frame in the spliced isoform. Retained introns/cassette exons and their flanking regions had to fit into the 162/147 nucleotide long variable region. As alternatively spliced introns and exons tend to be short, this did not constitute a severe limitation for the design.

#### Design of individual subsets

For each of the subsets in the libraries, a set of contexts from the previously assembled pool (as described above) was chosen based on the requirements of the specific question to be addressed, e.g. regarding the properties of the intron/alternative exon (length etc), complexity of the design scheme and required statistical power. Design of subsets was carried out in Python.

Multiple barcode controls: For each library we selected 25–40 splice site sequences expected to span a large range of splicing ratios and generated at least eight variants with identical variable region, but different barcodes.

Splice site mutations: For replacing the immediate splice site sequence with consensus and nonspliceable sequences, the following sequences were introduced: Donor splice sites: consensus (−3:+6) – CAGGTAAGT, nonspliceable (0:+6) – CTGCTC, GC-consensus (−3:+6) – CAGGCAAGT, U12-AT (0:+9) – ATATCCTTT, U12-GT (0:+9) – GTATCCTTT. Acceptor splice sites: consensus (– 15:+3) – CTCCTTTCCTTTCAGGTC, U12-acceptor (−19:+3) – TTCCTTAACTTCCTTTCAGATC, branch point (−26:−21) – CTCAC

Splice site switching/duplication: For 58 and 43 contexts containing tandem 5’ and 3’ splice sites, the immediate splice site sequences of variable length (9, 6 and 3 nt on the exonic side or increasing sequence portions (in increments of 3 nt) on the intronic side, up to the distance between the two splice sites) from either the first or the second splice site were used to replace the endogenous sequence in the respective other, leading to identical sequences upstream or downstream of the two splice sites.

Splicing factor binding sites: Binding sites for SRSF1 (TCACACGAC), SRSF2 (TGGCCTCTG), SRSF5 (TTCACAGGC), SRSF6 (CTGCGTCGA), hnRNPA1 (TTAGGGAAC), hnRNPG (CAAGTGTTC) and hnRNPU (TTGTATTGC), based on reports in the literature and experimental considerations (avoiding stop codons, restriction sites and homopolymers) were introduced at −58:−49, −49:−40, −40:−31, +5:+14, +9:+18 or +14:+23 relative to an acceptor and at −30:−21, −21:−12, −14:−5/−12:−3, +3:+12/+4:+13/+6:+15 or +12:+21/+13:+22/+15:+24 relative to a donor site, depending on the specific requirements imposed by the experimental design. Pairwise combinations of SRSF1, SRSF5, hnRNPA1 and hnRNPU were introduced to assay functional interactions, keeping a minimal distance of 9 nt between binding sites. Sequence motifs identified in previous studies were introduced into the same positions. Specifically, the sequences were CGACGTCGA, CAGAAGAGT, CGAAGATGT, CGCAAGAGT (“enhancers”), CCCAGCAGT, CCTTTTAGT, CCTAGTAGT (“silencers”), CAAAGAGGT, CAAACTTGT, CAACCTTGT (“neutral”), based on Ke et al. (2011) and adapted to accommodate the above mentioned experimental considerations. Hexamers (GENsil (“general silencing”): GTGGGG, E5enh (“enhancer in the alternative exon between tandem 5’ splice site “): CACCGC, E5sil (“silencer in the alternative exon between tandem 5’ splice site”): GGTGGG, I5enh (“enhancer in the intron downstream of tandem 5’ splice sites”): TTGTTC, I5sil (“silencer in the intron downstream of tandem 5’ splice sites”): CGAACC, E3enh (“enhancer in the alternative exon between tandem 3’ splice site “): CGAAGA, E3sil (“silencer in the alternative exon between tandem 3’ splice site”): GGGGGG, I3enh (“enhancer in the intron upstream of tandem 3’ splice sites”): TCTAAC, I3sil (“silencer in the intron upstream of tandem 3’ splice sites”): CCAAGC, identified by Rosenberg et al. (2015) were introduced into the same positions as above, with the three splice site-proximal positions in the 9 nt windows left unchanged.

Secondary structure: For changing local secondary structures around splice sites, two insertion sites per splice site were defined (with a length of 9 nt, introduced in frame such that the 3 nt upstream and 15 nt downstream of donor splice sites and 28 nt upstream and 3 nt downstream of acceptor splice sites were not changed). There, either the complement or the reverse complement for sequences at least 3 nt away (to allow for hairpin formation) were introduced, specifically −24:−15, 0:+9 and +3:+12 for donor splice sites and −9:0, −12:−3, +16:+25/+17:+26 for acceptor splice sites, depending on the specific requirements imposed by the experimental design.

Recoding and CG/GC: Most native sequence contexts were recoded either by random choice of synonymous codons or selection of synonymous codons with the highest or lowest GC content. The following triplets in frame were left unchanged so to not interfere with the basic functionality of the splice site: Eleven triplets before and one triplet after an acceptor site, as well as two full triplets (at least 6 nt, depending on the coding frame) before and after a donor site. CG and GC dinucleotides were introduced at different frequencies, leaving the 28 nt before and at least 3 nt after an acceptor as well as at least 3 nt before and at least 7 nt after a donor site unchanged.

Combinatorial variants: For 3–5 sets of contexts with equal intron or exon length or identical distance between two tandem splice sites (on average around 6 contexts in each set), all possible combinations of the three exonic, intronic or alternatively used exonic elements were created.

Introns and exons of identical length with no evidence for alternative splicing in RNAseq data (Encode, polyA RNA-seq of K562, Gingeras Lab, accession number ENCFF000HFA) were used to replace components of the alternative splice site contexts. For retained introns, all possible combinations of upstream exon, intron and downstream exon in all 38 contexts were replaced with the corresponding sequences from on average 3 constitutively spliced introns and their surrounding exons. For 5 groups of cassette exons of identical length, with around 6 contexts in each group, all components and combinations thereof where replaced with 5–6 constitutive exons and their surrounding intronic regions. In the case of tandem 5’ and 3’ splice sites, 8 exon-intron and 8 intron-exon regions with no evidence for alternative donor or acceptor sites were used to replace the corresponding exonic and intronic parts in 51 and 48 sequence contexts, respectively.

### Experimental procedures

#### K562 cell culture

K562 cells were acquired from ATCC. Cells were grown in Iscove’s Modified Dulbecco Medium supplemented with 10% fetal bovine serum (SIGMA) and 1% Penicillin-Streptomycin solution (SIGMA). The cells were split when reaching a concentration of ~10^6^ cells/ml. The cells were grown in an incubator at 37^o^C and 5% CO2. Cells were frozen in batches of 4×10^6^ cells in growth medium supplemented with 5% DMSO.

#### Construction of the master plasmid

Master plasmids for library insertion were constructed by amplifying parts from the genomic DNA or already existing vectors and cloning the parts sequentially into pZDonor 3.1. The master plasmid for the retained intron library contained the EF1alpha promoter, mCherry, a designed multiple cloning site containing restriction sites for library cloning (RsrII and AscI) and for inserting a downstream fragment (XbaI), GFP and the SV40 terminator sequence. As the alternative 5’ splice sites library only contained donor sites, the 3’ end of the intron (149 nt) and the beginning of the downstream exon (100 nt) corresponding to the EIF2D context used in the library were amplified from K562 genomic DNA (using primers tttccaGGCGCGCCtctgagagtggactgagtttggtt, tttccaTCTAGAgtcctggtaagtgtggagca) and cloned downstream of the library insertion site using AscI/XbaI. For the cloning of the cassette exon library, the 3’ end of the intron (722 nt) and the beginning of the exon (102 nt) downstream to a cassette exon in MCL1 were amplified from K562 genomic DNA (using primers tttccaGGCGCGCCttggagtggaagtagaatgaaggattt, tttccaTCTAGAtgccaaaccagctcctactccag) and cloned downstream of the library insertion site using AscI/XbaI.

#### Synthetic library cloning

The cloning steps were performed essentially as described previously (Vainberg Slutskin et al., 2018). We used Agilent oligo library synthesis technology to produce a pool of 55,000 different fully designed single-stranded 210-oligomers (Agilent Technologies, Santa Clara, CA), which was provided as a single pool of oligonucleotides (10 pmol). The four subsets of this pool corresponding to the libraries tested here were defined by unique amplification primers. The pool of oligos was dissolved in 200 μl Tris-ethylenediaminetetraacetic acid (Tris-EDTA) and then diluted 1:50 with Tris-EDTA, which was used as template for PCR. We amplified each of the four libraries by performing 8 PCR reactions, each of which contained 19 μl of water, 5 μl of DNA, 10 μl of 5× Herculase II reaction buffer, 5 μl of 2.5 mM deoxynucleotide triphosphate (dNTPs) each, 5 μl of 10 μM forward primer, 5 μl of 10 μM reverse primer, and 1 μl Herculase II fusion DNA polymerase (Agilent Technologies). The parameters for PCR were 95°C for 1 min, 14 cycles of 95°C for 20 s, and 68°C for 1 min, each, and finally one cycle of 68°C for 4 min. The oligonucleotides were amplified using library-specific common primers in the length of 35 nt, which have 18-nt complementary sequence to the single-stranded 210-mers and a tail of 17 nt containing RsrII (forward primer) and AscI (reverse primer) restriction sites. The PCR products were concentrated using Amicon Ultra, 0.5 ml 30K centrifugal filters (Merck Millipore). The concentrated DNA was then purified using a PCR mini-elute purification kit (Qiagen) according to the manufacturer’s protocol. Purified library DNA (540 ng total) was cut with the unique restriction enzymes RsrII and AscI (Fermentas FastDigest) for 2 hours at 37°C in two 40-μl reactions containing 4 μl fast digest (FD) buffer, 1 μl RsrII enzyme, 1 μl AscI enzyme, 18 μl DNA (15 ng/μl) and 16 μl water, followed by heat inactivation for 20 min at 65°C. Digested DNA was separated from smaller fragments and uncut PCR products by electrophoresis on a 2.5% agarose gel stained with GelStar (Cambrex Bio Science Rockland). Fragments were cut from the gel and eluted using electroelution Midi GeBAflex tubes (GeBA, Kfar Hanagid, Israel). Eluted DNA was precipitated using sodium acetate-isopropanol. The master plasmids were cut with RsrII and AscI (Fermentas FastDigest) in a reaction mixture containing 6 μl FD buffer, 3 μl of each enzyme and 3.5 μg of the plasmid in a total volume of 60 μl. After incubation for 2.5 hours at 37°C, 3 μl FD buffer, 3 μl alkaline phosphatase (Fermentas) and 24 μl water were added and the reactions were incubated for an additional 30 mins at 37°C followed by 20 min at 65°C. Digested DNA was purified using a PCR purification kit (Qiagen). The digested plasmids and DNA library were ligated for 30 min at room temperature in a 10 μl reactions, containing 150 ng plasmid and the library in a molar ratio of 1:1, 1 μl CloneDirect 10× ligation buffer, and 1 μl CloneSmart DNA ligase (Lucigen Corporation), followed by heat inactivation for 15 min at 70°C. Ligated DNA was transformed into *E. coli* 10G electrocompetent cells (Lucigen) divided into aliquots (23 μl each, plus 2 μl of the ligation mix), which were then plated on 4 Luria broth (LB) agar (200 mg/ml amp) 15-cm plates per transformation reaction (25 μl). For each library between 2 and 4 transformation reactions were performed. We collected between 0.5×10^6^ and 1.5×10^6^ colonies per library the day after transformation by scraping the plates into LB medium. Library-pooled plasmids were purified using a NucleoBond Xtra maxi kit (Macherey Nagel). To ensure that the collected plasmids contain only a single insert of the right size, we performed colony PCR (at least 16 random colonies per single transformation reaction).

For alternative 3’ splice sites and cassette exons, a common upstream donor site had to be introduced. To enable unambiguous identification of the variants based on the barcode at the 5’ end of the variable region, this had to be carried out after cloning of the library, as otherwise the 5’ end of the inserted library variants would be located in an intron and undetectable on the level of spliced mRNAs. In the case of alternative 3’ splice sites, the upstream exon (38 nt) and the 5’ end of the intron (132 nt) corresponding to the STAT3 context used in the library were amplified from K562 genomic DNA (using primers tttccaACTAGTgccccatacctgaagaccaag, tttccaGACGTCgtcactttgagtactaaacatagccc) and cloned into the library using AscI/XbaI, following the same protocol as above for the cloning of the oligonucleotide libraries. For the cloning of the cassette exon library, the upstream exon (52 nt) and the 5’ end of the intron (224 nt) upstream of the cassette exon in MCL1 from which the downstream sequences had been taken were amplified from K562 genomic DNA (using primers tttccaACTAGTttacgacgggttggggatgg, tttccaGACGTCttgacatcccaccctttccg) and cloned into the library using AscI/XbaI (see Fig S1A).

#### Transfection into K562 cells and genomic integration

The purified plasmid library was transfected into K562 cells and genomically integrated using the Zinc Finger Nuclease (ZFN) system for site-specific integration and the CompoZr^®^ Targeted Integration Kit - AAVS1 (SIGMA). Transfections were carried out using Amaxa^®^ Cell Line Nucleofector^®^ Kit V (LONZA). To ensure library representation we performed 10 nucleofections of the purified plasmid library. For each nucleofection, 4×10^6^ cells were centrifuged and washed twice with 20 ml of Hank’s Balanced Salt Solution (HBSS, SIGMA). Cells were resuspended in 100 μl solution (warmed to room temperature) composed of 82 μl solution V and 19 μl supplement (Amaxa^®^ Cell Line Nucleofector^^®^^ Kit V). Next, the cells were mixed with 2.75 μg of donor plasmid and 0.6 μg ZFN mRNA (prepared in-house) just prior to transfection. Nucleofection was carried out using program T-16 on the Nucleofector™ device, immediately mixed with ~0.5 ml of precultured growth medium and transferred to a 6 well plate with additional 1.5 ml of pre-cultured growth medium. A purified plasmid library was also transfected without the addition of ZFN and served as a control to determine when cells lost non-integrated plasmids.

#### Sorting the library by FACS

K562 cells were grown for at least 14 days to ensure that non-integrated plasmid DNA was eliminated. A day prior to sorting, cells were split to ~0.25×10^6^ cells/ml. On the day of sorting, cells were centrifuged, resuspended in sterile PBS and filtered using cell-strainer capped tubes (Becton Dickinson (BD) Falcon). Sorting was performed with BD FACSAria II SORP (special-order research product) at low sample flow rate and a sorting speed of ~18000 cells/s. To sort cells that integrated the reporter construct successfully and in a single copy (~4% of the population), we determined a gate according to mCherry fluorescence so that only mCherry-expressing cells corresponding to a single copy of the construct were sorted (mCherry single population). We collected a total of 3.1–3.9×10^6^ cells for each library (around 350 cells/variant on average) in order to ensure adequate library representation.

In the case of the retained introns library, cells sorted for single integration of the transgene were grown for a week before we sorted the population into 16 bins according to the GFP/mCherry ratio. Each bin was defined to span a range of GFP/mCherry ratio values such that it contains between 1–10% of the cell population. We collected a total of 1.2×10^7^ cells in order to ensure adequate library representation (>1000 cells/variant on average). Cells from each bin were grown separately for freezing and purification of genomic DNA.

#### RNA purification, cDNA synthesis, amplification and sample preparation

For the cell population sorted for single integration of the reporter construct we performed RNA purification by centrifuging 10^7^ cells, washing them with PBS, splitting into two tubes and purifying RNA using NucleoSpin RNA II kit (MACHEREY-NAGEL) according to the manufacturer’s protocol. We prepared cDNA in 4 reverse transcription reaction for each replicate using SuperScript^®^ III First-Strand Synthesis System (Thermo Fisher Scientific) with random hexamer primers and 5 μg of input RNA (per reaction) according to the manufacturer protocol. For amplification of the library variants, 3 PCR reactions of 50 ul total volume were performed. Each reaction contained 5 ul cDNA, 25 μl of Kapa Hifi ready mix X2 (KAPA Biosystems), 2.5 μl 10 μM 5’ primer, and 2.5 μl 10 μM 3’ primer. The PCR program was 95°C for 5 min, 20 cycles of 94°C for 30s and 72°C for 30s, each, and one cycle of 72°C for 5min. Specific primers corresponding to the constant region upstream and downstream of the splice sites were used. The PCR products were separated from potential unspecific fragments by electrophoresis on a 1.5% agarose gel stained with EtBr, cut from the gel, and cleaned in 2 steps: gel extraction kit (Qiagen) and SPRI beads (Agencourt AMPure XP). The sample was assessed for size and purity at the Tapestation, using high sensitivity D1K screenTape (Agilent Technologies, Santa Clara, California). We used 20 ng library DNA for library preparation for NGS; specific Illumina adaptors were added, and DNA was amplified using 14 amplification cycles. The sample was reanalyzed using Tapestation.

#### Genomic DNA purification, amplification and sample preparation

For each of the 16 bins of the retained intron library we purified genomic DNA by centrifuging 5×10^6^ cells, washing them with 1 ml PBS and purifying DNA using DNeasy Blood & Tissue Kit (Qiagen) according to the manufacturer’s protocol. In order to maintain the complexity of the library amplified from gDNA, PCR reactions were carried out on a gDNA amount calculated to contain a minimum average of 200 copies of each oligo included in the sample. For each of the 16 bins, we used 15 μg of gDNA as template in a two-step nested PCR. In the first step, three reactions were performed and each reaction contained 5 μg gDNA, 25 μl Kapa Hifi ready mix X2 (KAPA Biosystems), 2.5 μl 10 μM 5’ primer, and 2.5 μl of 10 μM 3’ primer. The parameters for the first PCR were 95°C for 5 min, 18 cycles of 94°C for 30s, 65°C for 30s, and 72°C for 60s, each, and one cycle of 72°C for 5min. In the second PCR step, each reaction contained 2.5 μl of the first PCR product, 25 μl of Kapa Hifi ready mix X2 (KAPA Biosystems), 2.5 μl 10 μM 5’ primer, and 2.5 μl 10 μM 3’ primer. The PCR program was 95°C for 5 min, 24 cycles of 94°C for 30s and 72°C for 30s, each, and one cycle of 72°C for 5min. Specific primers corresponding to the constant region of the plasmid were used. The 5’ primer contained a unique upstream 8-nt bin barcode sequence, and three different barcodes were used for each bin. The 3’ primer was common to all bins. Multiple PCR reaction products of each bin were combined. The concentration of the PCR samples was measured using a monochromator (Tecan i-control), and the samples were mixed in ratios corresponding to their ratio in the population, as defined when sorting the cells into the 16 bins. Sample preparation including gel elution and purification were performed as described above for amplicons from cDNA.

### Computational analyses

#### Mapping next generation sequencing reads

To unambiguously identify the variant of origin, a unique 12-mer barcode sequence was placed at the 5’ end of each variable region. DNA was sequenced on a NextSeq-500 sequencer. For cDNA we obtained 14.5, 4.3, 13.3 and 40.1 million reads for the retained intron, cassette exon, tandem 5’ and tandem 3’ libraries, respectively (2x150PE). Reads were first assigned according to their barcode (read 1) and subsequently the exact position of splicing (or lack thereof) was mapped using the corresponding mate (read 2) and assigned to either of the splice variants or, in the case of even a single mismatch or usage of a cryptic splice site, discarded. Both steps were performed using custom-made Python scripts.

For amplicons from genomic DNA from the 16 bins, into which the retained intron library was sorted, we obtained a total of ~12 million paired end reads (2x150bp, in order to cover the entire length of the variable region, not only the barcode, to filter out mutations introduced during synthesis or cloning, which could distort the protein readout (especially in the case of nonsense mutations and indels). Using Python scripts we determined for each read its bin barcode and its variant barcode and discarded all the reads that could not be assigned to a bin and a library variant of origin or contained even a single mismatch anywhere along the full length of the variant.

#### Computing RNA splicing ratios

For all variants with at least 100 reads mapped we computed the log2 ratio of spliced/unspliced reads for retained introns, exon included/exon skipped for cassette exons and downstream splice site used/upstream splice site used for tandem 5’ and 3’ splice site libraries, and refer to this throughout the text and figures as “splicing ratio” (in log2). A splicing ratio of 0 therefore indicates an equal number of reads mapping to the two possible splicing outcomes, with a positive value indicating more reads mapping to the spliced/”exon included”/”downstream splice site used” isoform and a negative value indicating more reads mapping unspliced/”exon skipped”/”upstream splice site used” isoform. In cases where more than 100 reads were mapped to a given variant, but all of them represented the same isoform, we added one read to the count of either isoform in order to enable us to calculate the log ratio for these variants. We chose to present splicing ratio as the ratio between the two expected outcomes, as opposed to PSI (percent spliced in) in order to have a measure that is meaningful across all splicing types tested here. In addition the log ratio between the two splice variants results in a larger dynamic range close to extreme values (0 or 100% spliced-in, i.e. dominance of one isoform). The small variance within barcode control groups around these values shows that our assay indeed is quantitative enough to draw conclusions even in this range.

After filtering we obtained RNA splicing ratios for 6626 (77.5%), 7249 (75.4%), 5266 (70.5%) and 4828 (67.5%) of the variants for the four libraries, respectively.

To determine normalized splicing ratios (i.e. the paired difference of a variant to the corresponding wild type context) we first calculated a mean splicing value (or noise value) for each context from triplicates (with different barcodes) added for all of the wild-type contexts. We then subtracted the corresponding mean wild-type level from each of the variants’ splicing values.

#### Computing protein splicing values

We applied a number of filters to the raw sequencing data to reduce experimental noise. First, variants with less than 200 reads mapped across bins were removed. Second, for bins with a read count of less than five or bins that got less than 2% of overall reads, the bin value was set to zero. Third, for each variant we set to zero bins surrounded by zero values (isolated bins). Forth, for each variant we set all cells to zero if the sum of normalized reads after filtering was less than 30% of the sum of normalized reads before filtering. For each variant, we normalized the values across the 16 bins and applied a Savitzky-Golay filter for smoothing the data. We detected peaks in the smoothed vector by a simple approach in which a point is considered a maximum peak if it has the maximal value, and was preceded (to the left) by a value lower by delta (which we set to 0.05). Variants with no or more than one peak after smoothing were disregarded in all protein-based analyses.

For each bin, we calculated the median of the log2 of GFP/mCherry as measured by FACS for all the cells sorted into that bin. For each variant, we calculated the weighted average and the variance for the distribution of reads across bins (using unsmoothed read counts normalized for each variant and taking the median of GFP/mCherry ratios of cells sorted into one bin as the value associated with this bin), resulting in what is referred to in the main text and figures as the “splicing value” (in log2) and, by dividing the variance by the mean, the “noise strength”. After filtering we obtained protein-based splicing values for 73% of the variants from our library of retained introns and 56% of the variants from our library of tandem 5’ splice sites. Noise residuals were calculated by fitting a linear equation to the relationship between noise strength and splicing value using scipy.stats.linregress and calculating the deviation of each point from this line.

#### Machine learning approaches

All machine learning procedures were carried out using the python sklearn package (version 0.18.2). Initially, from all duplicated sequences (e.g. barcode control sets), which passed filtering, a single variant was randomly chosen for all subsequent steps to avoid biases resulting from having duplicated sequences. 10% of the remaining variants were put aside and used only for evaluation of models built using the other 90%. We chose Gradient Boosting Regression as the prediction algorithm because it can capture non-linear interactions between features, which is especially relevant in the case of a complex problem like splicing prediction with many positional and combinatorial effects known.

For prediction based on hexamers, we counted the number of occurrences of every possible hexamer separately in the upstream exon, intron and downstream exon for retained introns, the upstream intron, exon and downstream intron for cassette exons and the exon, alternative exon and intron for tandem 5’ and 3’ splice sites, restricting ourselves to the designed variable region and disregarding the barcode.

For prediction based on hexamers, we used position weight matrices of RBP binding sites from the ATtRACT database (Giudice et al., 2016) to calculate the sum of log-odds ratios for all potential binding sites separately in the upstream exon, intron and downstream exon for retained introns, the upstream intron, exon and downstream intron for cassette exons and the exon, alternative exon and intron for tandem 5’ and 3’ splice sites, restricting ourselves to the designed variable region and disregarding the barcode.

For secondary structure predictions we used the fold function from the Vienna RNA package 2.0 and extracted both the minimal free energy and the predicted pairedness for each position.

Different hyperparameter settings for learning rate, n_estimators and max_depth were tested in a systematic and combinatorial fashion using 10-fold crossvalidation. Typically around 100 tests were performed and the best set of hyperparameters used for subsequent steps.

Feature selection was performed using optimized hyperparameters and sklearn’s feature_selection.SelectFromModel function. Another hyperparameter optimization step was performed to ensure that the previously chosen hyperparameters were still optimal for the reduced set of features.

At the end, the model was evaluated by training it on the entire training set (90% of all relevant unique library variants) and scoring the accuracy of prediction based on the held-out test set (10% of relevant unique library variants), which had not been used at any stage during development of the model. The R^2^ (coefficient of determination) regression score was chosen as a measure and calculated using the sklearn function metrics.r2_score.

#### General data analysis

For data analysis, we used python 2.7.11 with pandas 0.20.3, numpy 1.13.1, seaborn 0.6, scipy 0.17, and sklearn 0.18.2. Confidence intervals were calculated by bootstrapping (1000 iterations).

## Acknowledgements

The authors thank Adina Weinberger, Orna Dahan and members of the Segal and Pilpel labs for helpful discussions, Ronit Nir, Tali Avnit-Sagi and Maya Lotan-Pompan for technical advice and Shira Weingarten-Gabbay, Ronit Nir, Ilya Vainberg Slutskin and Tom Moss for critical reading of the manuscript. This work was supported by NIH and ERC grants (to E.S.) and an EMBO long-term fellowship (to M.M.).

## Author contributions

Conceptualization: M.M., Y.P. and E.S.; Methodology, Software and Formal Analysis: M.M.; Investigation: M.M. and A.H.; Writing – Original Draft: M.M.; Writing – Review & Editing: M.M. and E.S.; Funding Acquisition: M.M. and E.S.; Supervision: Y.P. and E.S.

The authors declare no competing interests.

## References

Amit, M., Donyo, M., Hollander, D., Goren, A., Kim, E., Gelfman, S., Lev-Maor, G., Burstein, D., Schwartz, S., Postolsky, B., et al. (2012). Differential GC Content between Exons and Introns Establishes Distinct Strategies of Splice-Site Recognition. Cell Rep. 1, 543–556.

Änkö, M.-L. (2014). Regulation of gene expression programmes by serine-arginine rich splicing factors. Semin. Cell Dev. Biol. 32, 11–21.

Barash, Y., Calarco, J.A., Gao, W., Pan, Q., Wang, X., Shai, O., Blencowe, B.J., and Frey, B.J. (2010). Deciphering the splicing code. Nature 465, 53–59.

Braunschweig, U., Gueroussov, S., Plocik, A., Graveley, B.R., and Blencowe, B.J. (2013). Dynamic integration of splicing within gene regulatory pathways. Cell 152, 1252–1269.

Cieply, B., and Carstens, R.P. (2015). Functional roles of alternative splicing factors in human disease. Wiley Interdiscip. Rev. RNA 6, 311–326.

Eldar, A., and Elowitz, M.B. (2010). Functional roles for noise in genetic circuits. Nature 467, 167–173.

Elowitz, M.B., Levine, A.J., Siggia, E.D., and Swain, P.S. (2002). Stochastic gene expression in a single cell. Science 297, 1183–1186.

Giudice, G., Sánchez-Cabo, F., Torroja, C., and Lara-Pezzi, E. (2016). ATtRACT— a database of RNA-binding proteins and associated motifs. Database J. Biol. Databases Curation 2016.

Gurskaya, N.G., Staroverov, D.B., Zhang, L., Fradkov, A.F., Markina, N.M., Pereverzev, A.P., and Lukyanov, K.A. (2012). Analysis of alternative splicing of cassette exons at single-cell level using two fluorescent proteins. Nucleic Acids Res. 40, e57.

Hicks, M.J., Mueller, W.F., Shepard, P.J., and Hertel, K.J. (2010). Competing Upstream 5ʹ Splice Sites Enhance the Rate of Proximal Splicing. Mol. Cell. Biol. 30, 1878–1886.

Jangi, M., and Sharp, P.A. (2014). BUILDING ROBUST TRANSCRIPTOMES WITH MASTER SPLICING FACTORS. Cell 159, 487–498.

Kaufmann, B.B., and van Oudenaarden, A. (2007). Stochastic gene expression: from single molecules to the proteome. Curr. Opin. Genet. Dev. 17, 107–112.

Ke, S., Shang, S., Kalachikov, S.M., Morozova, I., Yu, L., Russo, J.J., Ju, J., and Chasin, L.A. (2011). Quantitative evaluation of all hexamers as exonic splicing elements. Genome Res. 21, 1360–1374.

Ke, S., Anquetil, V., Zamalloa, J.R., Maity, A., Yang, A., Arias, M.A., Kalachikov, S., Russo, J.J., Ju, J., and Chasin, L.A. (2018). Saturation mutagenesis reveals manifold determinants of exon definition. Genome Res. 28, 11–24.

Kim, D., Shivakumar, M., Han, S., Sinclair, M.S., Lee, Y.-J., Zheng, Y., Olopade, O.I., Kim, D., and Lee, Y. (2018). Population-dependent Intron Retention and DNA Methylation in Breast Cancer. Mol. Cancer Res.

Lev Maor, G., Yearim, A., and Ast, G. (2015). The alternative role of DNA methylation in splicing regulation. Trends Genet. 31, 274–280.

Marinov, G.K., Williams, B.A., McCue, K., Schroth, G.P., Gertz, J., Myers, R.M., and Wold, B.J. (2014). From single-cell to cell-pool transcriptomes: stochasticity in gene expression and RNA splicing. Genome Res. 24, 496–510.

McManus, C.J., and Graveley, B.R. (2011). RNA structure and the mechanisms of alternative splicing. Curr. Opin. Genet. Dev. 21, 373–379.

Ozbudak, E.M., Thattai, M., Kurtser, I., Grossman, A.D., and van Oudenaarden, A. (2002). Regulation of noise in the expression of a single gene. Nat. Genet. 31, 69–73.

Rosenberg, A.B., Patwardhan, R.P., Shendure, J., and Seelig, G. (2015). Learning the Sequence Determinants of Alternative Splicing from Millions of Random Sequences. Cell 163, 698–711.

Shalek, A.K., Satija, R., Adiconis, X., Gertner, R.S., Gaublomme, J.T., Raychowdhury, R., Schwartz, S., Yosef, N., Malboeuf, C., Lu, D., et al. (2013). Single-cell transcriptomics reveals bimodality in expression and splicing in immune cells. Nature 498, 236–240.

Vainberg Slutskin, I., Weingarten-Gabbay, S., Nir, R., Weinberger, A., and Segal, E. (2018). Unraveling the determinants of microRNA mediated regulation using a massively parallel reporter assay. Nat. Commun. 9.

Waks, Z., Klein, A.M., and Silver, P.A. (2011). Cell‐to‐cell variability of alternative RNA splicing. Mol. Syst. Biol. 7, 506.

Wong, J.J.-L., Gao, D., Nguyen, T.V., Kwok, C.-T., van Geldermalsen, M., Middleton, R., Pinello, N., Thoeng, A., Nagarajah, R., Holst, J., et al. (2017). Intron retention is regulated by altered MeCP2-mediated splicing factor recruitment. Nat. Commun. 8, 15134.

Xiong, H.Y., Alipanahi, B., Lee, L.J., Bretschneider, H., Merico, D., Yuen, R.K.C., Hua, Y., Gueroussov, S., Najafabadi, H.S., Hughes, T.R., et al. (2015). The human splicing code reveals new insights into the genetic determinants of disease. Science 347, 1254806.

Yeo, G., and Burge, C.B. (2004). Maximum entropy modeling of short sequence motifs with applications to RNA splicing signals. J. Comput. Biol. J. Comput. Mol. Cell Biol. 11, 377–394.

